# Paradigm shift in biomarker translation: a pipeline to generate clinical grade biomarker candidates from DIA-MS discovery

**DOI:** 10.1101/2024.03.20.586018

**Authors:** Qin Fu, Manasa Vegesna, Niveda Sundararaman, Eugen Damoc, Tabiwang N. Arrey, Anna Pashkova, Emebet Mengesha, Philip Debbas, Sandy Joung, Dalin Li, Susan Cheng, Jonathan Braun, Dermot P.B. McGovern, Christopher Murray, Yue Xuan, Jennifer E. Van Eyk

## Abstract

Clinical biomarker development has been stymied by inaccurate protein quantification from mass spectrometry (MS) discovery data and a prolonged validation process. To mitigate these issues, we created the Targeted Extraction Assessment of Quantification (TEAQ) software package. This innovative tool uses the discovery cohort analysis to select precursors, peptides, and proteins that adhere to established targeted assay criteria. TEAQ was applied to Data-Independent Acquisition MS data from plasma samples acquired on an Orbitrap™ Astral™ MS. Identified precursors were evaluated for linearity, specificity, repeatability, reproducibility, and intra-protein correlation from 11-point loading curves under three throughputs, to develop a resource for clinical-grade targeted assays. From a clinical cohort of individuals with inflammatory bowel disease (n=492), TEAQ successfully identified 1116 signature peptides for 327 quantifiable proteins from 1180 identified proteins. Embedding stringent selection criteria adaptable to targeted assay development into the analysis of discovery data will streamline the transition to validation and clinical studies.

## Introduction

Clinical proteomics for the development and translation of disease biomarkers typically involve three phases: discovery, targeted protein assay development and clinical validation^1-6^. In the initial discovery phase, high-throughput mass spectrometry (MS) screens are commonly employed to identify biomarker candidates within a defined disease cohort^7-9^. Targeted protein assays are then individually developed by manually determining reliably quantifiable representative peptide(s) for each biomarker candidate. These assays are used to validate the biomarkers in large, more diverse clinical cohorts. The progression of biomarker candidates from discovery through the validation stages can be a time-consuming process, often spanning several years^10,11^. To date, this process has yielded less than desirable success translating candidate biomarkers to clinical use. Two challenges that impede the transition include peak intensity data that cannot be reliably quantified for a significant fraction of the identifiable proteins and the time-intensive development of targeted protein assays that meet the analytical performance needed for large scale validation studies. A clear avenue to improve this pipeline is increasing the analytical rigor applied in the discovery phase to improve selection of candidates worthy of further validation.

In the discovery phase, the data independent acquisition (DIA)-MS for analyzing plasma samples offers several advantages^12^ over traditional data dependent acquisition (DDA)-MS methods. Unlike DDA-MS, which selects precursor ions for fragmentation based on their intensity^13^. DIA-MS systematically fragments all precursors within a series of predefined mass windows. Plasma samples are dominated by a small number of highly abundant proteins that can bias DDA-MS acquisitions. DIA-MS’s comprehensive approach increases plasma proteome coverage and reduces missing values, making it the preferred method for complex biological samples in clinical proteomics^14-16^.

Following candidate identification, targeted protein approaches such as Multiple Reaction Monitoring (MRM) with triple quadrupole mass spectrometers or Parallel Reaction Monitoring (PRM) with high-resolution mass spectrometers^17-20^ are employed to develop assays that monitor signature peptides representative of the candidate biomarker^21,22^. Signature peptides are selected by empirically determining their correlation with the candidate protein as well as characterizing their linear response, reproducibility, lower limit of detection (LLOD) and quantitation (LLOQ) in the matrix^23,24^. The resulting assays are quantified with stable isotopic labeled (SIL) standards to provide greater sensitivity and accuracy compared to discovery methods. However, targeted approaches have their limitations, including requiring prior knowledge of the target protein and difficulty in high content multiplexing^25,26^. We have identified an advantage in embedding targeted assay development concepts into the discovery phase. Though, this requires achieving the same level of precision, repeatability, reproducibly, linear range, *etc*. characteristic of the dedicated targeted approaches.

The Orbitrap Astral MS platform is a high-resolution accurate mass (HRAM) instrument, which integrates advanced mass analyzers^27^ to produce high quality mass spectra very rapidly across a wide dynamic range of peptide concentrations^27-29^. Here we evaluate the Orbitrap Astral MS across various parameters including linearity, specificity, repeatability, with the goal of evaluating the quantifiability of DIA data and embedding dedicated targeted techniques in the discovery phase. Initially, we evaluated an 8-point dilution curve of a pooled human plasma sample with three different sample throughput methods. To leverage the Orbitrap Astral MS’s advanced performance and facilitate targeted assay development, we developed TEAQ (Targeted Extraction Assessment of Quantification), a software package which automates the selection of prototypic signature peptides that meet rigorous standards for reliability, reproducibility, linearity, and precision from DIA-MS data. Using the Orbitrap Astral MS, we demonstrate that it is possible to identify signature peptides from DIA-MS data. These signature peptide data sets, provide a global resource for selecting the appropriate throughput, for discovery and can be bench marked using the 53 FDA or LDT approved protein markers present in the data set.

We further tested this approach for biomarker development by applying our workflow to an inflammatory bowel diseases (IBD) and control cohorts (n=492). IBD represents a spectrum of chronic, hyper-inflammatory diseases primarily involving the gut^30-32^. IBD is a lifelong disease, with a peak onset in teenage and early adult years, associated with poor quality of life affecting the ability to work, and a need for recurrent surgeries^33^. There is a clear unmet need for IBD biomarkers that can diagnose, monitor disease activity, measure therapeutic effectiveness and predict recurrence^34,35^.

Our stringent approach to discovery DIA-MS uses the Orbitrap Astral MS platform and TEAQ identified signature peptides to meet the technical requirements for clinical grade assays in biomarker validation. Collectively, we have provided a resource for facilitating targeted assay development as well as the TEAQ tool for deriving signature peptides from new cohorts and experiments.

## Results

### Orbitrap Astral MS DIA throughput and performance

In this study, we evaluated the performance of the Neo-Vanquish™ HPLC Orbitrap Astral MS platform using a pooled digested human plasma sample by DIA-MS across three different throughputs: 180 samples per day (SPD) with a 8-min runtime, 100 SPD with a 14-min runtime, and 60 SPD with a 24-min runtime (n=5). We also compared our findings with previously published data obtained using an Orbitrap Exploris™ 480 MS coupled with the Evosep One (60 SPD) using the same plasma (Figure 1A, 1B and 1C). As anticipated, longer acquisition runtimes (lower SPD) yielded more identified and quantifiable proteins. The Orbitrap Astral MS exhibited excellent performance across all sample throughput protocols. In our analysis, 88-91% of precursors were observed in all 5 injections with median CV of precursor intensities ranging from 6% to 17%. For example, using the 180 SPD, 568 proteins (1% FDR) were identified in all injections and 358 of those further met the CV<20% criteria. At 60 SPD, we identified a total of 649 proteins with 1% FDR. Remarkably, 89% of the proteins (576) were observed in all 5 technical replicates and 78% of these plasma proteins (504) had a CV<20%, meeting the standard for clinical-grade assays.

**Figure 1.**
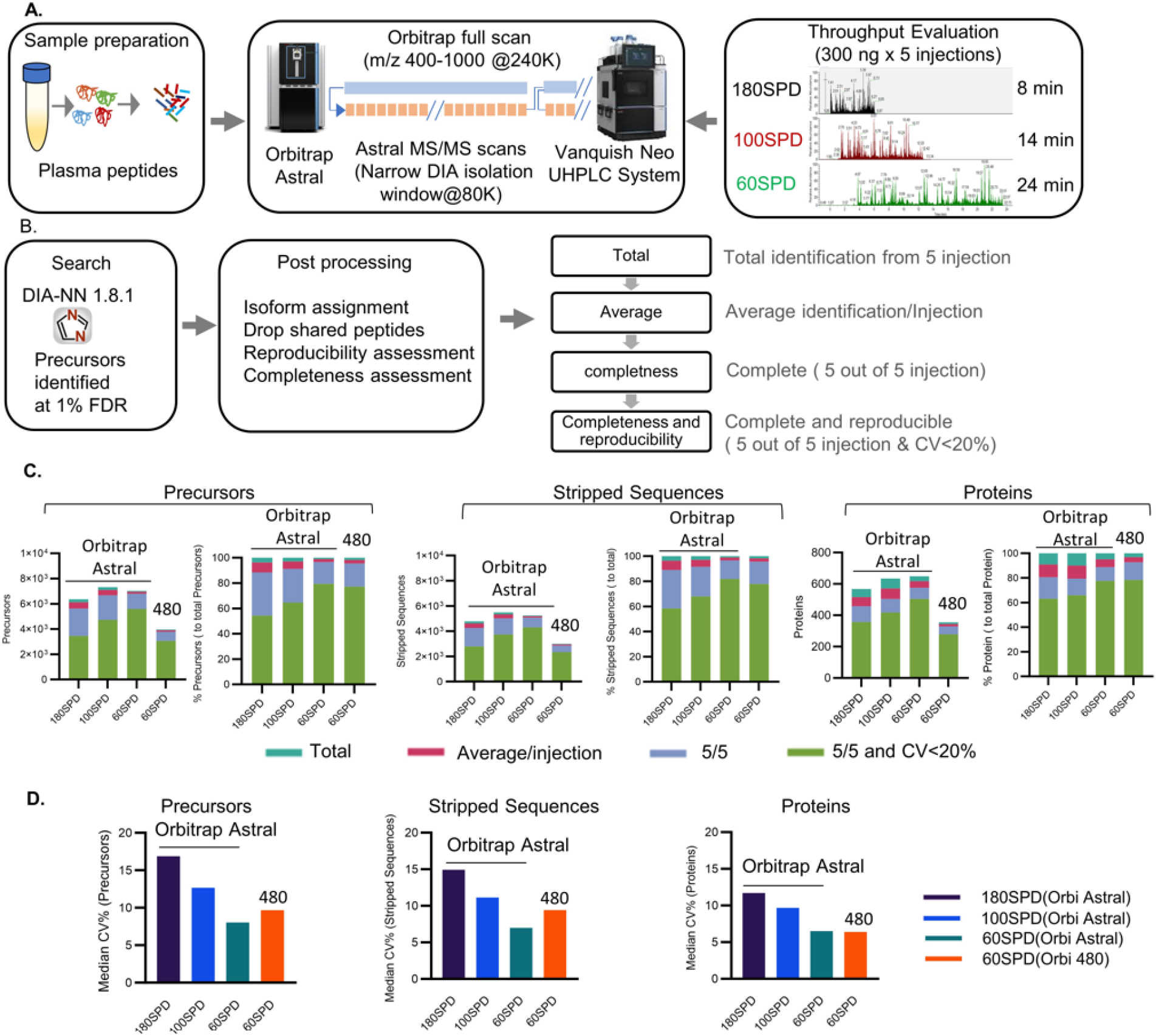
The workflow to evaluate Orbitrap Astral MS performance for plasma proteome. **A**. Representation of parallelized acquisition DIA method, with Orbitrap full MS scans overlapped with Astral MS/MS scans. Liquid chromatograph system was a Vanquish Neo UHPLC system operated in trap-and-elute configuration. Three different throughputs were evaluated, digested plasma samples were run with 24 min (60 samples per day or SPD), 14 minutes (100SPD) or 8 minutes (180SPD) total runtime with PepMapTM columns. Human plasma K2EDTA gender pooled plasma from100 healthy females and healthy 100 males were purchased commercially. **B**. Data analysis pipeline. Raw files were analyzed in DIA-NN 1.8.1 with an in-silico DIA-NN predicted spectral library with iRT. The output results from DIA-NN (precursor at 1% FDR) were analyzed. The shared precursors (peptides) among different proteins were dropped and precursors shared among isoform rollup to canonical proteins. **C**. Bench marker of Orbitrap Astral in combination of Neo-Venquish LCMS/MS with three different throughputs, 180SPD, 100SPD and 60SPD with 300 ng on column loading. The identified total precursors from repeated injections (n=5), average precursors per injection, precursors with data completeness in repeated injection (n=5), and with precision (CV<20%) were calculated. For comparison, Orbitrap Exploris 480 60SPD throughput with 250 ng on column loading were also included in the performance evaluation. **D**. Evaluate precision for the three Orbitrap Astral methods and Orbitrap Exploris 480. Median CV% were calculated from 5 injections of 300 ng on column for Orbitrap Astral and 250 ng on column for Orbitrap Exploris 480 were calculated.

Compared to the previously published Exploris 480 MS data set on the same pooled plasma sample, the Orbitrap Astral MS demonstrated superior sensitivity, reproducibility, and completeness (Figure 1C)^36^, identifying 1.7-fold more proteins with a lower median CV (Figure 1C). Notably, 97% of precursors were detected in five out of five runs and 80% were detected on all five runs with a CV<20% (Figure 1C). Our results with 300 ng loading demonstrated that 60 SPD had the best completeness and reproducibility of the throughput methods tested (Figure1D).

Next, we evaluated plasma precursors suitable for potential targeted assay development using a loading curve assay. Digested pooled plasma was serially diluted to create an 8-point curve ranging from 12.5ng to 500ng loaded on column. Each point was injected 5 times for each of the previously established throughput methods (180, 100 and 60 SPD). Search results were evaluated for total precursor identifications, reproducibility, and data completeness (Figure 2A). 100 and 60 SDP methods yielded an average of 10% more precursors than the 180 SPD method. Of the precursors identified, the vast majority were observed in 5 out of 5 injections across all the points in the loading curve demonstrating high data completeness. The precursor measurements also exhibited good reproducibility with a mean CV<20% for each of the methods for columns loading of 200ng and higher. When considering the combined parameters of completeness (5 out of 5 observations) and reproducibility (CV<20%), the different throughput methods exhibited a distinct hierarchy: 60SPD > 100SPD > 180SPD.

**Figure 2.**
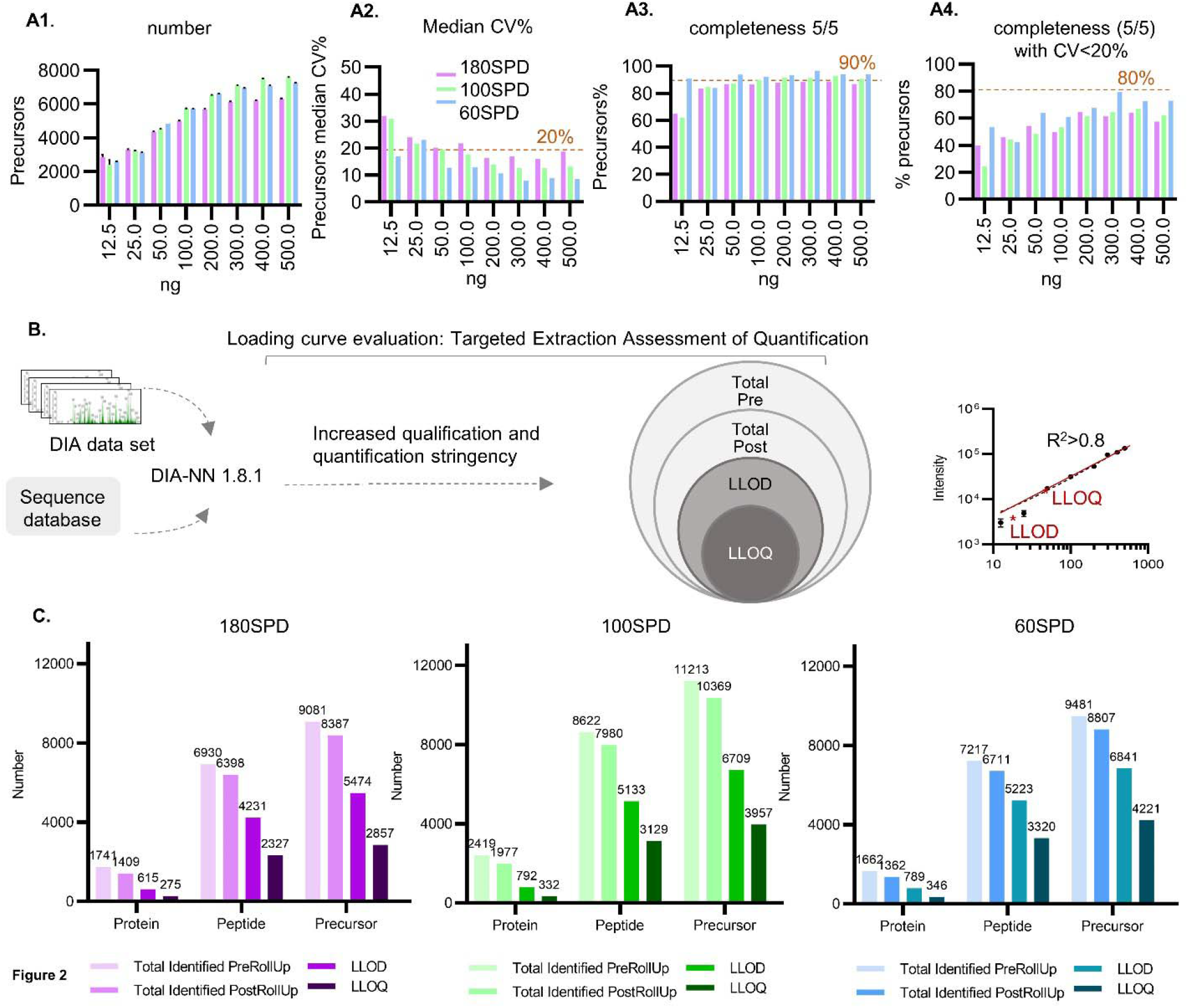
Development of TEAQ pipeline and three throughputs evaluation via loading curves. **A**. Comparative Analysis of Precursor. The evaluation involves comparing the total number of identified precursors, the number of precursors with LLOD, and the number of precursors with LLOQ across three throughputs: 180SPD, 100SPD, and 60SPD. Each loading curve has eight loading points (5 repeated injections, 12.5 ng to 500 ng). **A1**, The average total precursor. **A2**. Median Coefficient of Variation (CV%). **A3**. The percentage of completeness. A4. The percentage of precursors observed in all 5 injections and CV<20%. **B**. The TEAQ process evaluates both the quality and quantifiability of the DIA dataset. Raw files underwent an initial conversion to the mzML format and were searched using DIA-NN 1.8.1, employing a library-free approach. The DIA datasets initially underwent a quality assessment, focusing on reproducibility and completeness. Subsequently, the datasets underwent targeted extraction with assessment of quantification (TEAQ). This involved evaluating various parameters: including total identified precursors; reliably identified (lowest loading with >60% completeness or present in 3/5 injections); those with a lower limit of detection (LLOD, defined as the lowest loading with >60% completeness and CV<20%), those with a lower limit of quantification (LLOQ, defined as the lowest concentration with linear regression R2 >0.8, CV<20% and recovery 80%-120%). **C**. Proteins, peptides, and precursors (total identified both before and after rollups, LLOD and LLOQ), were detected across the entire 5x8 loading points curve for 180SPD, 100SPD, and 60SPD throughputs.

### Development of TEAQ workflow for DIA analysis: Loading curve study

The initial TEAQ analysis uses loading curve data to categorize identified precursors into a series of progressively stringent groupings (Figure 2B). After total identifications, TEAQ grouped all the reliably observed species (present in 60% of injections). From that list an LLOD was determined based on the lowest point in the curve with a CV<20%. The list was further refined to determine an LLOQ, where the R^2^>0.8 for 3 consecutive points and the lowest point had a CV<20% and a target deviation <20%. This analysis was conducted for each throughput method (Figure 2C). Using the 60 SPD method, TEAQ identified a total of 8807 precursors representing 1362 proteins, of which 789 and 346 had characterizable LLODs and LLOQs, respectively. Similar results were found for the other two throughput methods. The percentage of identified precursors where an LLOQ was determined was 31% (180SPD), 35% (100SPD) and 45% (60SPD), demonstrating the positive correlation between longer gradients and identification of quantifiable analytes. Detailed loading curve results for the three throughput methods, including individual precursor evaluations, are presented in Supplemental tables 1-3.

### Development of TEAQ workflow for DIA analysis: clinical cohort study

To assess the suitability of our Orbitrap Astral MS-TEAQ pipeline for biomarker development, we evaluated plasma proteomes from 205 individuals diagnosed with IBD and 287 age, sex and race matched healthy control subjects (Supplemental Table 4).

Our laboratory has previously established a standardized workflow for achieving precise high-throughput proteomics analysis of blood biofluids, specifically plasma, employing an Biomek i7 automated liquid handling workstation (Figure 3A)^37^. For the discovery cohort analysis, we opted for the 60SPD configuration with a 24-min runtime. All 492 plasma samples were prepared using 6x96-well plates, following the published automated protocol^36^. Each plate required two days to acquire, and the entire study acquisition spanned approximately 12 days. Indexed retention time standards (iRTs) were spiked into each sample prior to MS analysis to monitor the longitudinal performance of the LC MS/MS across the entire acquisition.

**Figure 3.**
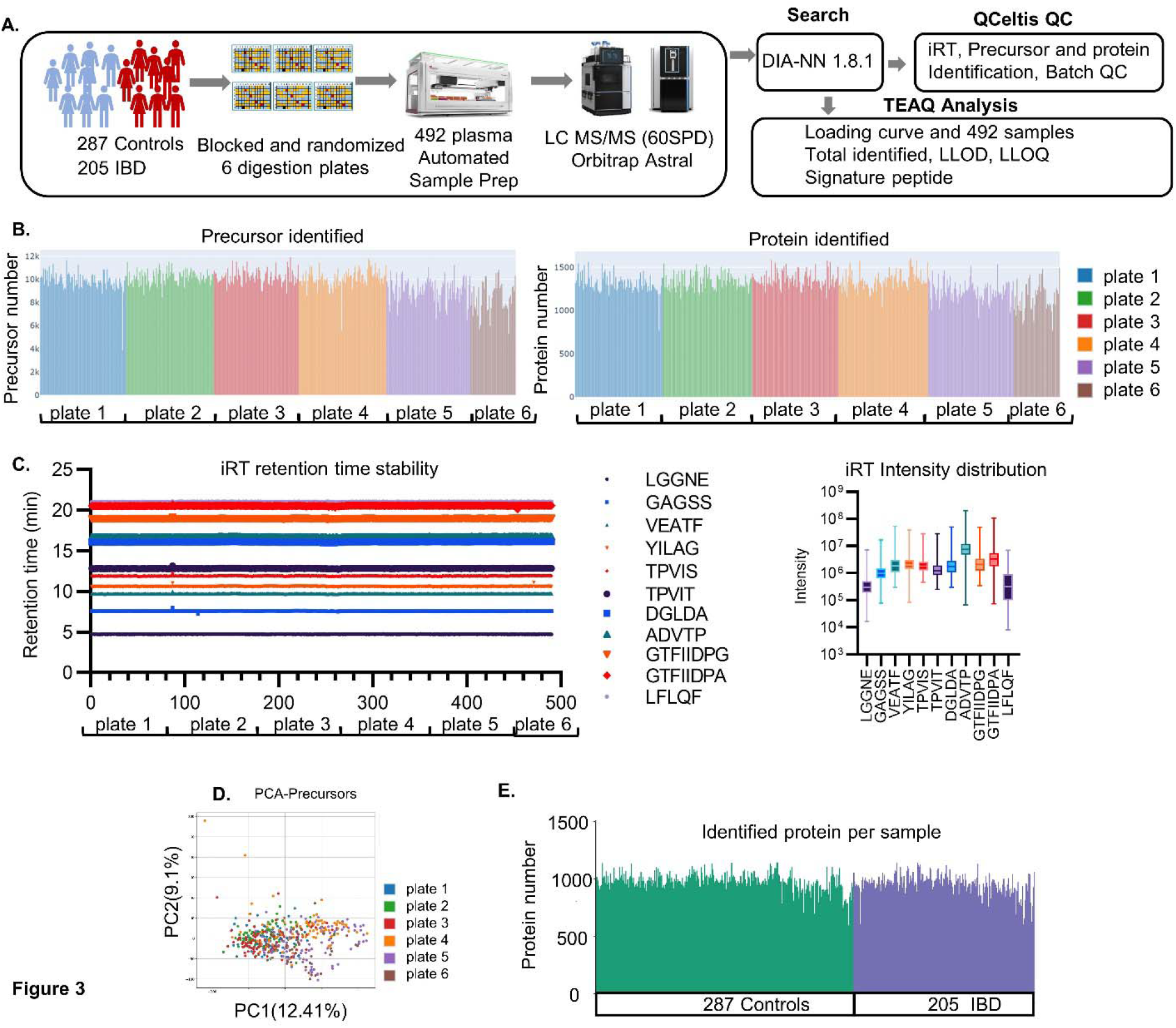
IBD cohort sample preparation, data search, QC and proteins identification. **A**. 492 IBD cohort samples (287 controls and 205 IBD subjects) randomized across 6 plates and processed with i7 automation workstation. Data acquisition was performed by A Vanquish Neo UHPLC system coupled to an Orbitrap Astral mass spectrometer with an Easy spray™ source. The raw data was first converted to .mzML and then search via DIANN 1.8.1 with a predicted library and precursors were identified with 1% FDR. Data quality control analysis on individual samples as well as batch (plate based) were performed by Qcelits. **B**. The identified pre-rollup precursors and proteins numbers were plotted batch/plate based. **C**. Spiked 11 standard reference peptides (iRT) from all 492 samples retention time and intensity were extracted and plotted. Each peptide sequence is denoted by the first five letters of its amino acid sequence. **D**. Principle analysis of precursor intensities from all 6 batches/plates. **E**. Average proteins per sample after protein isoform rollup and shared peptides removing in the control and IBD groups were plotted. Each colored line in the plot represents a single sample.

Data was searched using DIA-NN library free mode employing key quality parameters illustrated in Figure 3A. After extracting precursors, peptides, and proteins, we assessed the robustness and consistency of the identified proteins with our QCeltis quality control (QC) software package. The QCeltis package from our laboratory monitors metrics based on the input datasets and provides visual QC summary of a data set (manuscript in preparation). We observed an average of >9700 precursors and >1300 proteins pre rollup, per sample (Figure 3B). Post rollup, we found an average of 923±62 and 903±70 proteins per sample in the control and IBD groups, respectively (Figure 3E). The Vanquish Neo UHPLC system exhibited exceptional retention stability throughout the entire experiment, spanning 6 plates acquired over 12 days as illustrated by iRT (Figure 3C). Furthermore, Principal Component Analysis (PCA) of precursor intensities from the 6 plates (Figure 3D) demonstrated that all plates clustered together, suggesting that batch correction was not necessary. We identified a similar number of proteins in control and IBD groups (Figure 3E) indicating that missingness was not skewed to one group.

Next, we employed the TEAQ pipeline (Summarized in Supplemental Figure 1) to examine the protein expression profiles of the control and IBD subjects. We identified a total of 11,493 precursors, representing 8,655 stripped sequences, and 1,180 proteins observed in at least 20% of samples, post-rollup, per group (Supplemental Figure 2 and Supplemental Figure3). The TEAQ package further refined the precursors, stripped peptides, and proteins by removing miss-cleaved peptides and retaining only those unique to the entire FASTA database. TEAQ then applied the data from the loading curve to define 4,986 precursors (representing 3,819 peptides and 407 proteins) with a determined LLOQ. The TEAQ pipeline analysis of the IBD/control cohort samples, in combination with loading curve, resulted in a total of 1,225 precursors and 1,116 peptides, representing 327 proteins, classified as quantifiable with precision and correlated signature peptides (Supplemental Figure 3).

A signature peptide, as per our definition, exhibits a unique peptide sequence specific to the target protein, with no miscleavages, a linearity R^2^ >0.8, and a CV<20%. Additional criteria were that deviations <20% at the lower limit of quantification and in cases where multiple peptides or precursors originate from the same protein, that all peptides demonstrate a correlation coefficient within the same proteins (>0.7) in the study.

### Candidate IBD biomarkers

The Orbitrap Astral MS and TEAQ identified several candidate biomarkers from the IBD discovery cohort. We selected four proteins (CRP, A1AG1, COMP, and PHLD) as examples to illustrate the value of TEAQ and its automated analysis of pre and post rollup precursors, LLOD, LLOQ and the identification of signature precursors/peptides from DIA-MS IBD data set (Figure 4A). Notably, each protein has a unique count in each category; CRP, and COMP had at least one signature peptide that met all criteria. PHLD and A1AG1 had 9 and 5 correlated signature peptides, respectively. Among these four proteins, two are upregulated, and two are downregulated in IBD, with p-values ranging from 0.004 to <0.0001. For each protein, 1 to 2 signature peptide signal responses along the 11-point loading curves are presented, highlighting the excellent quantitation (R^2^, LLOQ) for each signature peptide(s) in each protein (Figure 4B).

**Figure 4:**
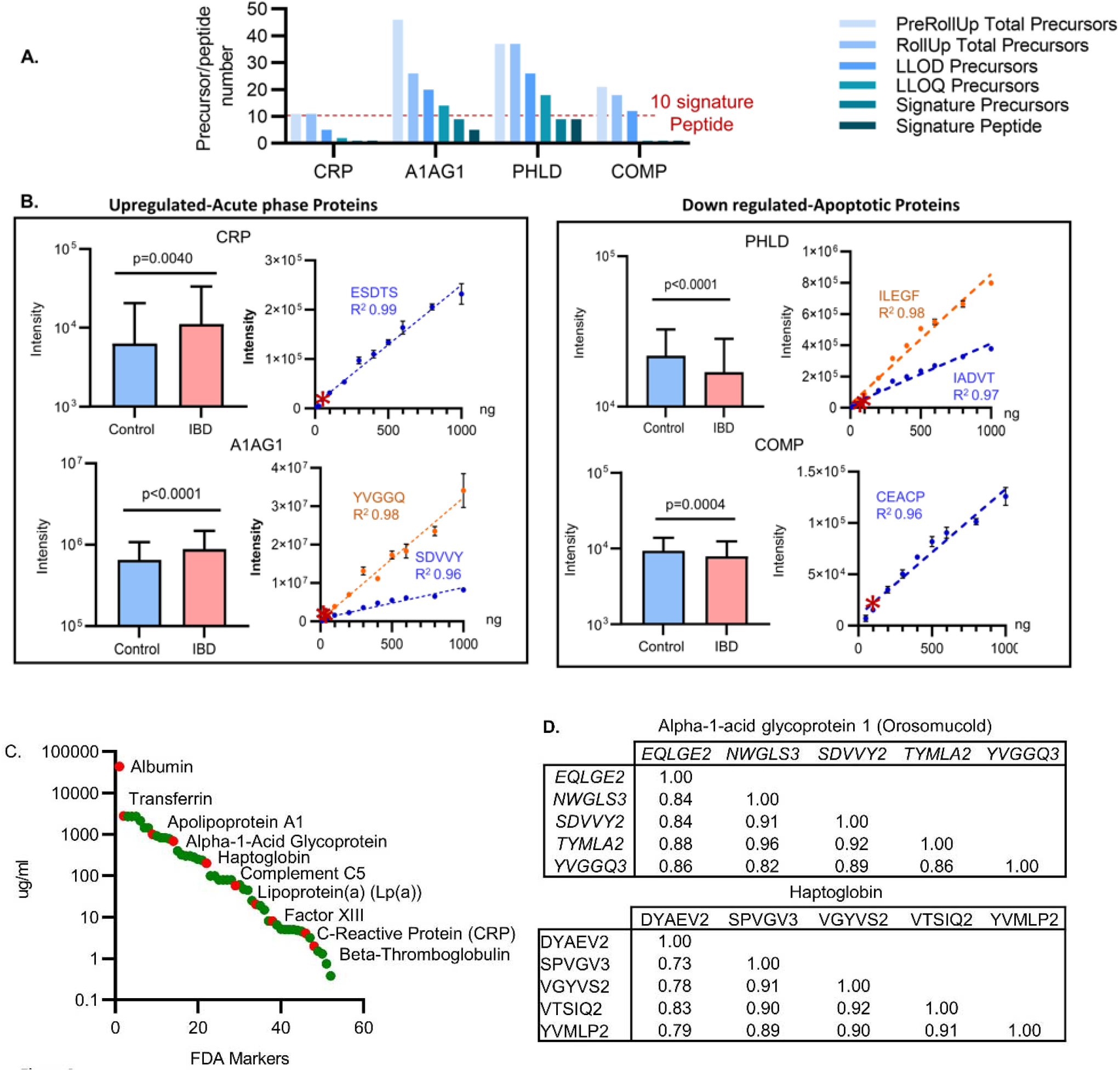
Selected 4 markers in IBD cohort and identified FDA markers with TEAQ pipeline. **A**. Four proteins were chosen as representative proteins from the TEAQ processing package. The pre-rollup total precursors, rollup total precursors, LLOD precursors, LLOQ precursors, signature precursors, and signature peptides observed from 492 samples (287 controls and 205 IBD subjects) are presented for CRP, A1AG1, PHLD and COMP. Statistical analysis was performed by Prism (version 9.5.1) employing an unpaired T-test, assuming a Gaussian distribution. Bar graphs depicting the intensity of IBD and control were generated by inputting protein intensities from individual samples (control n=287, IBD n=205) into Prism, which plotted the mean with standard deviation. For the calibration curve, an XY plot with error bars was generated by Prism using the entered “mean & CV%” values, and the R^2^ value was calculated by Prism. **B**. Two upregulated proteins (CRP and A1AG1) and two downregulated proteins (PHLD and COMP) are plotted, displaying their expression levels and corresponding p-values. The selected signature peptides, along with their R^2^ values and LLOQ (*), are plotted based on 5 injections x 11 loading points ranging from 12.5 ng to 1000 ng using a 60SPD workflow. **C**. FDA markers. 53 FDA markers found in IBD cohort from TEAQ package. **D**. Two examples showcase the correlation of signature peptide intensity for FDA proteins within a IBD cohort of 492 subjects. Alpha-1-acid glycoprotein 1 (also known as Orosomucold or AGP1) and Haptoglobin, both have five correlating signature peptides. Each peptide sequence is denoted by the first five letters of its amino acid sequence, followed by a numeric value indicating the charge state.

Of particular interest, two acute phase proteins (CRP and A1AG1) were upregulated with IBD. CRP, a general inflammation marker (FDA approved), has been reported to elevate in response to inflammation in IBD patients^38^. Currently, CRP is measured as part of diagnostic and monitoring procedures for IBD patients^39^. TEAQ selected a signature peptide (ESDTSYVSLK) that has been previously validated using internal standards and an antibody-based ELISA assay^40,41^. A1AG1, also known as orosomucoid protein, is known acute phase proteins and function in modulating the activity of the immune system^42^, were reported to be associated with IBD^43^ and age dependent^44^. In addition, A1AG1 (FDA approved) is one of four potentially useful circulating biomarkers for estimating the five-year risk of all-cause mortality (the other three are albumin, very low-density lipoprotein particle size, and citrate)^45^.

We have also highlighted two apoptotic proteins (COMP and PHLD) that were reduced in the plasma of IBD patients. PHLD (Glycosylphosphatidylinositol Specific Phospholipase D1) primarily from liver and COMP (Cartilage Oligomeric Matrix Protein) in number of different tissues, are all secreted proteins and involved in the apoptosis process. Their disparate origins suggest a body wide impact of the inflammatory condition. The specific associations between PHLD and COMP, and their role in apoptosis in the context of Inflammatory Bowel Disease (IBD) were not widely reported or well-established.

### General utility of our Orbitrap Astral Plasma signature precursor/peptide database

By performing an initial pooled loading curve experiment, TEAQ’s sequential procedure whittles down DIA results to only those that meet the stringent criteria necessary for targeted assay development. This initial characterization can be used to filter a discovery cohort analysis to select the most reliable precursors for validation. Our initial loading curve TEAQ analysis was performed using healthy pool plasma across sample throughput methods using the Orbitrap Astral MS. While performing loading curves on individual discovery cohorts is ideal, many potential biomarkers represent changes in concentration of proteins commonly present in healthy plasma negating the need for repeated loading curve analysis. For example, our 60 SPD loading curve analysis characterized a total of 1,225 signature precursors (1,116 peptides), representing 327 proteins (Supplemental Figure 3) with 53 currently FDA-approved markers. We offer the complete TEAQ loading curve analysis of the three sample throughput methods as a resource to expedite PRM/MRM targeted assay development (Supplemental Table 1, 2 and 3).

## Discussion

The three traditional phases of biomarker development are time-consuming and often fail to yield clinically viable biomarkers. Our study presents a novel paradigm applying assay development-level stringency to high-throughput discovery screening, facilitating the identification of mature, validation-ready biomarkers. Our aim is to prune poorly performing candidates in the discovery phase, accelerating clinical assay development and validation to expedite the translation of final biomarkers to clinical practice.

In the targeted protein assay development phase, the precision and accuracy of protein quantification relies entirely on the careful selection of suitable signature peptides, acting as surrogates for the protein of interest. The current approach is to experimentally test multiple candidate peptides and select those with the highest correlation^46^. In practical mass spectrometry-based targeted quantification approaches, not all peptides are observed and quantified equally. This discrepancy arises from various unpredictable factors, including ionization efficiency, amino acid composition, site-specific digestion efficiency, matrix effects, LC interference, and other unknown variables. The development of a true quantitative targeted assay often requires trial and error to identify reliably quantifiable signature peptides for a protein of interest. To meet clinical assay standards, signature peptides should be unique to the protein of interest, free of missed cleavages, and amenable to detection and quantification by LC-MS/MS. They should yield linear, reproducible signals in LC-MS/MS analysis, as evidenced by dose-response curves (reflecting the LLOQ). Additionally, each peptide should be represented by a distinct MS1 mass (precursor mass) and demonstrate correlated LC-MS/MS signals across multiple peptides (or precursors) within the study population. The painstaking selection and characterization of these peptides is a limiting factor in constructing highly multiplexed quantitative targeted assays for biomarker validation^22^.

To realize the potential of biomarker development there needs to be a robust pipeline including discovery-based LC-MS to allow for reliable technical data to be obtained. The Orbitrap Astral MS platform offers extensive proteome coverage and comprehensive data, identifying over 1,000 proteins from digested plasma regardless of SPD. TEAQ streamlines biomarker candidate screening using stringent quantification criteria, minimizing the bottleneck of a lengthy assay development (Figure 5). Supplemental Table 5 summarized the characteristics of TEAQ in comparison to Clinical Laboratory Standard institute (CLSI) LCMS based proteins and peptide assay development criteria (Quantitative Measurement of Proteins and peptides by Mass Spectrometery C64 1st edition (2021) by Clinical Laboratory Standards Institute).

**Figure 5.**
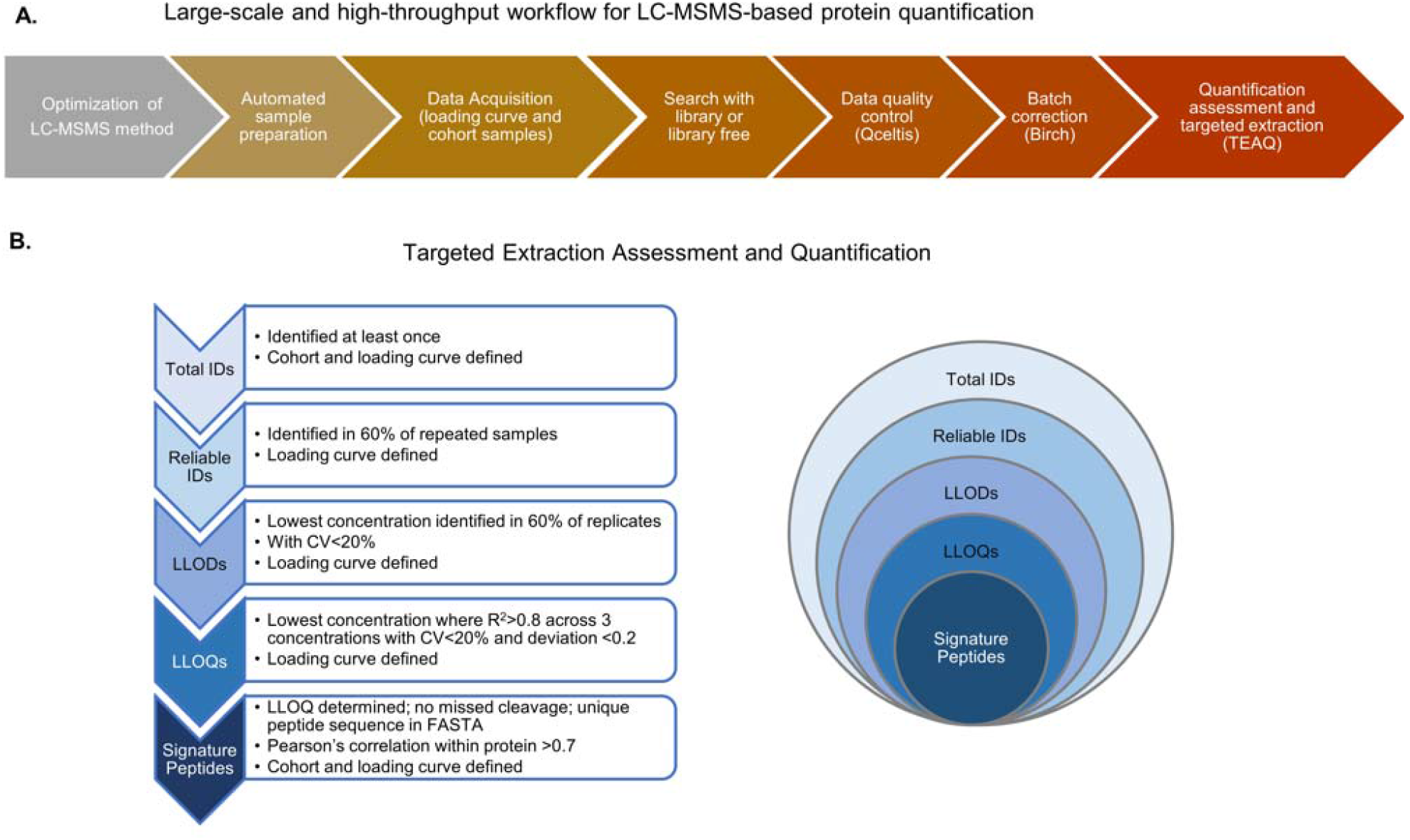
High throughput workflow and TEAQ. **A**. Multiple and sequential steps involved in LCMSMS based proteins quantification. Our solutions and software packages are indicated with parentheses. Loading curve with replicated injections and at least > 6 loading point. QCeltis (manuscript in preparation), Birch^47^ and TEAQ. **B**. The schema of Targeted Extraction Assessment of Quantification workflow (TEAQ) automated workflow.

Using the Orbitrap Astral MS and TEAQ, we identified candidate biomarkers from an IBD discovery cohort. Notably, we identified two acute-phase proteins upregulated in IBD: CRP and A1AG1, as well as two apoptotic proteins reduced in IBD patients: PHLD and COMP. While our TEAQ workflow has limitations, such as the lack of internal standards in the current IBD cohort, future enhancements may incorporate SIL information to enable more accurate quantification in subsequent analyses.

## Conclusion

Our automated and unbiased TEAQ pipeline groups DIA discovery data into total observed, reliably observed, LOD, LOQ, and signature peptides (precursors) with increased stringency. We implemented our pipeline to identify reliably quantifiable proteins within an inflammatory bowel disease (IBD) cohort. Fifty-three FDA or LDT proteins commonly used in clinical settings were quantified in our IBD cohort. Our methodology introduces a novel approach to biomarker research, bridging the gap between discovery and validation, thereby facilitating the translation of biomarkers to the clinic.

## Materials and Methods

### Plasma samples

Human plasma K_2_EDTA gender pooled from 100 healthy females and 100 healthy males (0.2μm filtered) purchased from BioIVT (Westbury, NY) was used for the 8-or 11-point concentration curves. The sample pooled plasma was used previously^36,37^.

Details of the IBD and control cohorts has been described previously^48,49^. In brief, in both cohorts’ study subjects were enrolled in an IRB-approved prospective registry at Cedars-Sinai Medical Center (CSMC) and plasma samples were collect between December 2020 and June 2021. The IBD cohort consists of subjects 13+ years old with IBD diagnosis based on standard clinical criteria, and the HCW cohort consists of healthcare professionals in CSMC. Both cohorts were initially designed to study the effect of SARS-COVID-2 vaccines and received two doses of mRNA vaccine as per local department of health guidelines and institutional policies, and the samples were collected 16 weeks after the 2nd dose of COVID vaccine in both cohorts. IBD cohort plasma samples consisted of IBD subjects (n=205) collected as outline in Kumar *et al*. ^50^ along with age and sex matched controls (n=287) from the relatively healthy health care worker cohort, Corale^51,52^. Supplemental table 4 lists the sex, age and race characteristics of IBDs and controls. All human plasma was aliquoted and stored at -80°C until processing. Clinical samples were prepared^36^, then analyzed with the optimized DIA method at a throughput of 60 sample per day (SPD).

### Automated plasma samples digestion and desalting

Automated plasma sample digestion and desalting were carried out as previously published^36^. Plasma samples were processed using the BioMek i7 automated workstation (Beckman Coulter)^37,53^. Briefly, plasma samples (5 μL) were denatured in 350 mL/L 2,2,2-trifluoroethanol (TFE), 40mM dithiothreitol in 50mM NH4CO3 at 60°C for one hour, and then alkylated in 10 mM iodoacetamide at 25°C for 30 min. Reactions were quenched with 5 mM dithiothreitol at 25°C for 15 min. Samples were diluted with 50mM NH_4_CO_3_ (final TFE concentration of 50 ml/L) and digested with trypsin (sample to enzyme ratio of 25:1) and incubated for 4 hours at 37°C. The digestion reaction was stopped by adding 20% formic acid to a final concentration of 2% formic acid. Desalting was carried out using a positive pressure apparatus (Amplius Positive Pressure ALP, Beckman Coulter) mounted on the left side of the i7 Biomek workstation deck^53^. Digested samples (300 μL) were mixed with 850 μl of 2% phosphoric acid, 0.1% formic acid and applied to an Oasis 30 µm HLB 96-well Plate (Waters Co.). After 1x ml MeOH activation and 3x 1ml 0.1% formic acid, 1150 μL samples were loaded on the HLB plate and eluted with 1 ml of 50% acetonitrile, 0.1% formic acid and evaporated to dryness and stored at -80□C until use. At the time of MS analysis, peptides were resuspended in a 0.1% formic acid solution and iRT was added prior to the LC MSMS run.

### DIA LC-MS/MS analysis

A Vanquish™ Neo UHPLC system (Thermo Fisher Scientific) coupled to an Orbitrap™ Astral™ MS with an EASY-spray™ source (Thermo Fisher Scientific) was used to acquire data. Digested plasma samples were separated using a 24 min (60 SPD), 14 min (100SPD) or 8 min (180SPD) total runtime with PepMap™ columns 150 μm x 15 cm (Thermo Fisher Scientific) in a trap&elute configuration. The mobile phases were 0.1% FA in water (A) and 0.1% FA in 100% ACN (B). For the Orbitrap Astral MS data acquisition, HRAM Orbitrap MS1 scans were collected in parallel with ultra-high speed (up to 200 Hz) HRAM Astral DIA MS2 scans. MS1 spectra were acquired at resolution of 240,000 with a mass range of 400 to 1000 m/z. Normalized automatic gain control (AGC) target value for fragment spectra of 500% was used. MS2 spectra were acquired with various narrow isolation widths and injection times (i.e., 2 Th / 3.5 ms or 3 Th / 5 ms). Peptide ions were fragmented at normalized collision energy (NCE) of 25%.

For loading curves, the digested peptide samples were analyzed in replicates (5 injections for each sample at each concentration). The loading curves were performed by serial dilution of digested and desalted human gender pooled plasma with 0.1% FA in water. The experiments were carried out with the optimized ultra-high throughput (180 SPD) and higher throughput (100SPD and 60SPD) methods for 8 different sample loads (12.5, 25, 50, 100, 200, 300, 400 and 500 ng on column). In addition, one 60SPD method with 5 ms injection time had an 11-point loading curve (12.5, 25, 50, 100, 200, 300, 400, 500, 600, 800 and 1000 ng on column). For IBD clinical cohort (n=492), a single injection of 60SPD method with 5ms injection time was used. To avoid carryovers, blank runs with Zebra-wash (4 Zebra wash cycles on trap column) were inserted every 20 sample.

The previously published Orbitrap Exploris 480 MS data set was used for comparison^36^.

### Bioinformatic Data Analysis

Loading curve raw files were converted to mzML using MSConvert (Version: 3.0.23229-ac2773e). For a loading curve, DIA-NN (DIA-NN software package, version 1.8.1)^54^ search was performed independently for each concentration using an in silico digested human reviewed and canonical FASTA library downloaded from the UniProt database (December 2020). Retention time (RT)-dependent cross run normalization was enabled. Library generation was done with enabling “FASTA digest for library-free search” and “Deep learning-based spectra, RTs and IMs prediction”, and Match Between Runs (MBR) was also enabled within each search (RTs= retention times and IMs= ion mobilities). When reporting protein numbers and quantities, the Protein.Ids column in DIA-NN’s main report was used to identify the protein group and the Precursor.Normalized column was used to obtain the normalized precursor quantities. The precursor m/z range was set between 400 and 1000, and the fragment ion m/z range was set between 200 and 1800. Additionally, the missed cleavages rate was set to 2 and peptide length range was set between 5 to 30. The protein inference parameter was set to “Protein Names”. All other settings were left default. The software output was filtered at precursor q-value <1% and the global protein q-value <1% filter was also applied to all benchmarks.

DIA-NN Search and reporting for IBD cohort samples was performed similarly except that all converted mzML files for 492 raw files were searched in one search were MBR and RT-dependent cross run normalization was enabled.

**QCeltis** QCeltis is a quality control analysis software package from our laboratory (manuscript in preparation). This package uses plots to view large amounts of information to identify patterns and trends across the samples and datasets. This python package was used for performing quality control analysis on the IBD cohort samples across all 6 plates. Protein and precursor-level intensity files from the DIA-NN search results and plate maps were provided as input to the QCeltis package. The enzyme parameter was set to ‘Trypsin’ and the default parameters values for TIC CV, CV Percent and Data Percent Thresholds were used. Additional thresholds were protein (300), precursor (3000) and missed cleavage. The QCeltis analysis produced Excel and HTML reports with the quality control metrics, visualizations, and results.

**TEAQ**. TEAQ, our in-house python package evaluates independent DIA-NN search results from the loading curve and cohort analysis and extract signature precursors for peptides that meet the criteria of for targeted clinical assay development (Supplemental Figure 1). A precursor represents an individual charge state observation of a peptide in MS1. Since a unique precursor mass is required to build PRM/MRM assays, TEAQ package outputs can be directly used in assay development. A signature precursor is defined as a peptide sequence that is unique to the target protein, exhibiting no miscleavages (presence of one K or one R at the C-terminal of a peptide), a linearity R^2^ > 0.8 and a coefficient of variation (CV%) below 20%. Additionally, the deviation of the LLOQ concentration should be less than 20%. In cases where multiple peptides/precursors originate from the same protein, it is essential that all peptides demonstrate correlation within the same proteins (>0.7) in the study.

Precursor-level data for the loading curve was first filtered to remove shared peptides/precursors among different “Protein IDs” column in the DIA-NN search output, thereby only retaining peptides assigning to unique canonical proteins (and isoforms). Additionally, for peptides shared across canonical proteins and their corresponding isoforms, a roll-up was done by choosing the canonical protein as the representative protein and summing intensities across its isoforms. Next, the lower limit of detection (LLOD) was determined for each precursor by selecting the lowest sample concentration where the normalized precursor intensity was detected in 3/5 replicates (observation value = 0.6) with a CV ≤20%. The lower limit of quantification (LLOQ) for each precursor was estimated as the lowest sample concentration satisfying the linearity criterion of square of correlation coefficient (R^2^) ≥0.8 for a minimum of three consecutive concentration points and a target deviation of less than 20%. Precursors meeting both the LLOD and LLOQ criteria were chosen as quantifiable precursors.

The list of quantifiable precursors was then applied to the IBD dataset to determine cohort specific signature precursors. The cohort dataset was filtered according to the following criteria: observations >20% across cohort samples, peptide sequences unique to FASTA database and had no missed cleavages, while precursors with charge +1 were removed if the same precursor is present with a higher charge state. Next, intensity values were used to calculate a Pearson’s correlation coefficient for each remaining precursor and compared to its parent protein. Only peptide/precursors that satisfy the correlation coefficient criteria (R^2^) ≥ 0.7. This remaining list of peptide/precursors represents the available signature peptides/precursors. Access to TEAQ is available by contacting the corresponding author.

## Supporting information

SupplementTable1_180SPD_8min-3p5ms_Linearity

SupplementTable2_100SPD_14min-3p5ms_Linearity

SupplementTable3_60SPD_24min-5ms_Linearity

SupplementTable4_CaseControlsCharacteristics

SupplementTable5_AssaydevelopmentTEAQandCLSI

## Supplemental Figures

**Supplemental Figure 1.**
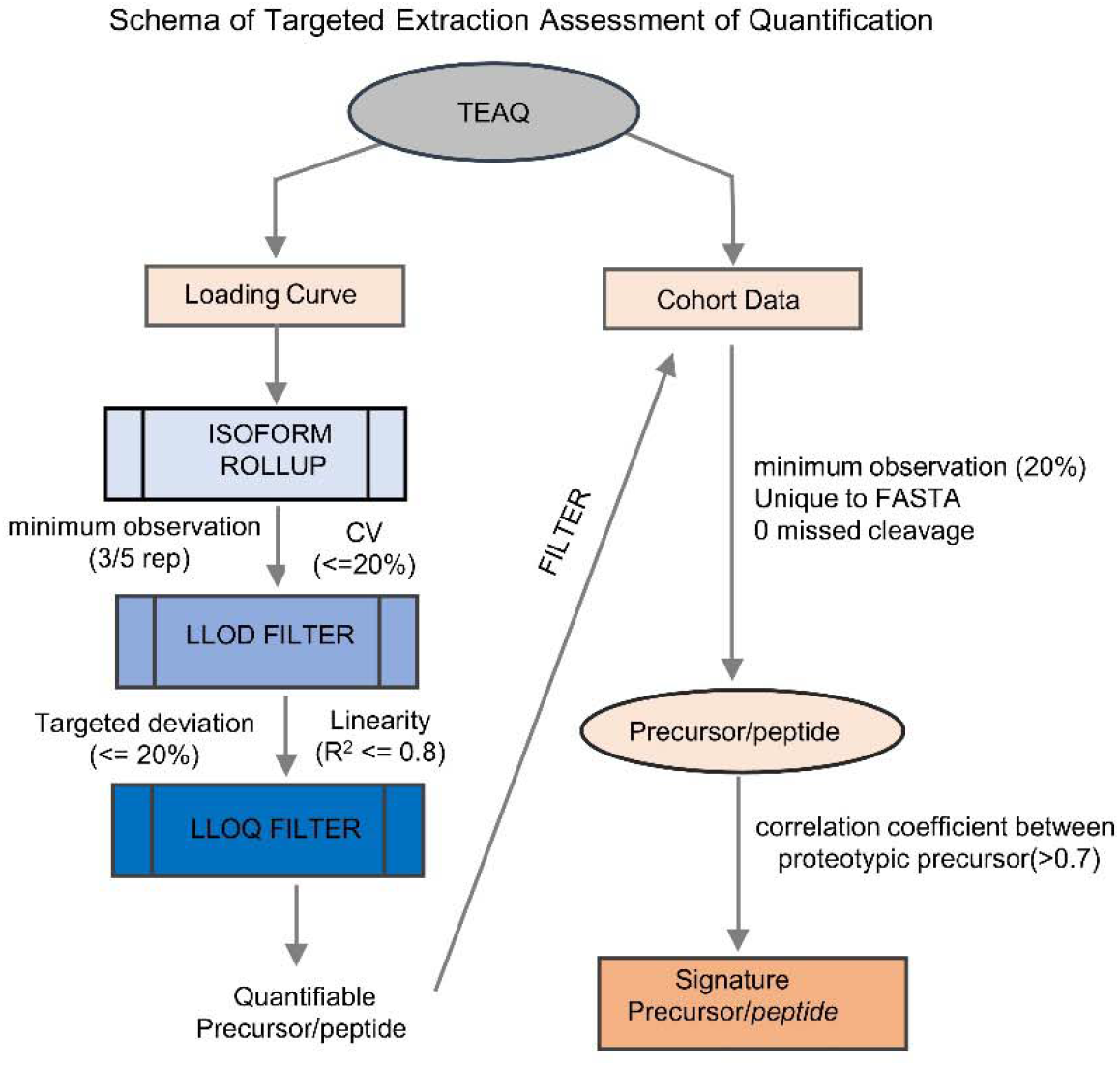
TEAQ workflow. TEAQ, a software package developed for performing targeted extraction of quality peptides or precursors to enhance the validation of biomarker discovery from large-scale proteomics datasets. It facilitates the selection of prototypic peptides and precursors that meet rigorous standards for reliability, reproducibility, linearity and precision from DIA datasets.

**Supplemental Figure 2.**
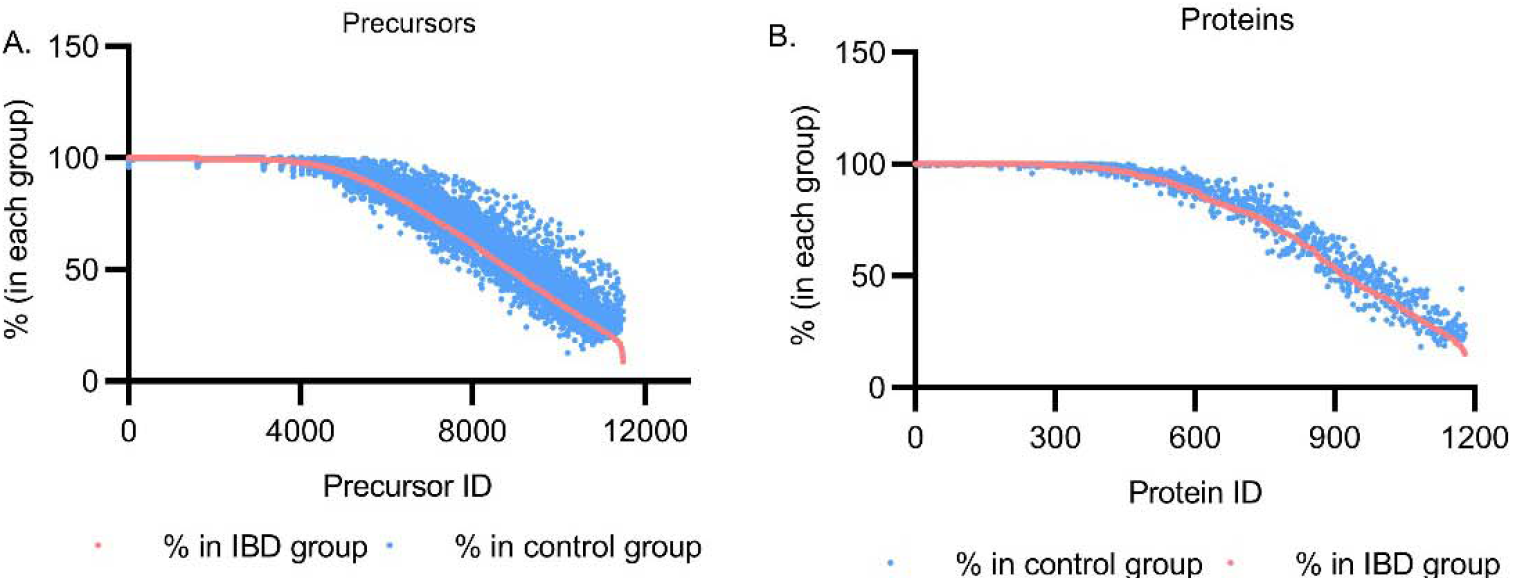
Precursors and proteins observed in the IBD group or in the control group. Precursors and proteins presented in IBD group or control group were examined. Each precursor’s presence was calculated as a percentage specific to either the IBD group or the control group. A. All identified precursors in the IBD group were found to also be present in the control group. B. All identified proteins the IBD group or in the control group.

**Supplemental Figure 3.**
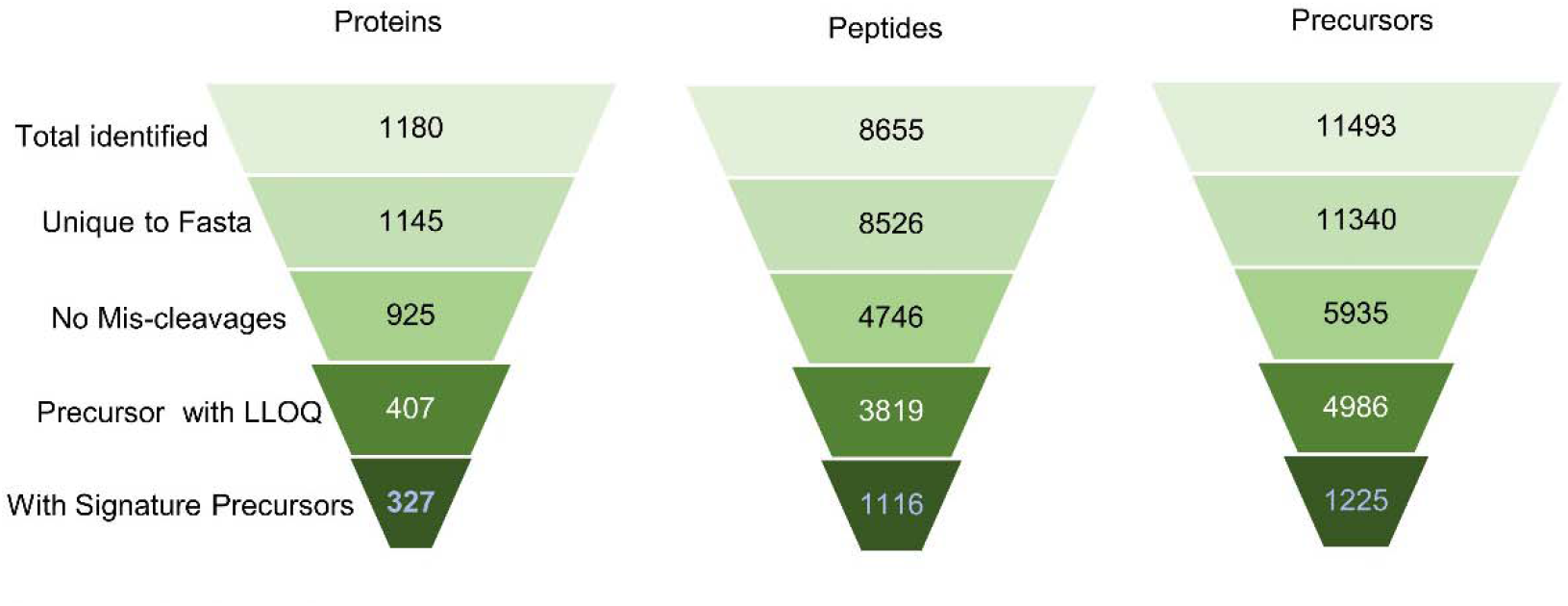
TEAQ and observed precursors, peptides and proteins in entire IBD study. From the entire IBD study cohort (n=487), the following observations were made regarding proteins, peptides, and precursors: total identified, unique to the FASTA database, absence of miss-cleavage, lower limit of quantification (LLOQ), and signature precursors (representing peptides and proteins) within each group.

### Ethics

Ethical approval was obtained from the Institutional Review Board on Research Involving Human Subjects (Cedars-Sinai Medical Center IRB#: Study00001411 and Study00000621).

## Acknowledgement

The authors wish to express their gratitude for the contributions that have greatly facilitated the success of this study. Special thanks are extended to Koen Raedschelders for his assistance in sample organization and aliquoting, Ali Haghani for his support in sample shipment, as well as Maxim Zhgamadze, Nathan Hendricks and Angel Keoseyan for sample preparation.

This study was supported by 1 U01 DK124019-01 (JVE), 1 R01 HL155346-01A1(JVE), the Leona M. and Harry B. Helmsley Charitable Trust, the Widjaja Foundation Inflammatory Bowel and Immunobiology Research Institute, National Institute of Diabetes and Digestive and Kidney Disease Grants P01DK046763 and U01DK062413, the Cedars-Sinai Precision Health Initiative, and the Erika J. Glazer Family Foundation.

## Notes

### Competing Interest Statement

1. YX, ED, TNA and AP are employees of Thermo Fisher Scientific
2. QF and JVE are inventors of US Patent 10,352,94

## References

1 Cilento, E. M. et al. Mass spectrometry: A platform for biomarker discovery and validation for Alzheimer’s and Parkinson’s diseases. J Neurochem 151, 397–416 (2019). 10.1111/jnc.14635

2 Nakayasu, E. S. et al. Tutorial: best practices and considerations for mass-spectrometry-based protein biomarker discovery and validation. Nat Protoc 16, 3737–3760 (2021). 10.1038/s41596-021-00566-6

3 Parker, C. E. & Borchers, C. H. Mass spectrometry based biomarker discovery, verification, and validation--quality assurance and control of protein biomarker assays. Mol Oncol 8, 840–858 (2014). 10.1016/j.molonc.2014.03.006

4 Macklin, A., Khan, S. & Kislinger, T. Recent advances in mass spectrometry based clinical proteomics: applications to cancer research. Clin Proteomics 17, 17 (2020). 10.1186/s12014-020-09283-w

5 Zhang, B. et al. Clinical potential of mass spectrometry-based proteogenomics. Nat Rev Clin Oncol 16, 256–268 (2019). 10.1038/s41571-018-0135-7

6 Birhanu, A. G. Mass spectrometry-based proteomics as an emerging tool in clinical laboratories. Clin Proteomics 20, 32 (2023). 10.1186/s12014-023-09424-x

7 Crutchfield, C. A., Thomas, S. N., Sokoll, L. J. & Chan, D. W. Advances in mass spectrometry-based clinical biomarker discovery. Clin Proteomics 13, 1 (2016). 10.1186/s12014-015-9102-9

8 Messner, C. B. et al. Ultra-High-Throughput Clinical Proteomics Reveals Classifiers of COVID-19 Infection. Cell Syst 11, 11–24 e14 (2020). 10.1016/j.cels.2020.05.012

9 Hawkridge, A. M. & Muddiman, D. C. Mass spectrometry-based biomarker discovery: toward a global proteome index of individuality. Annu Rev Anal Chem (Palo Alto Calif) 2, 265–277 (2009). 10.1146/annurev.anchem.1.031207.112942

10 Boja, E. S., Fehniger, T. E., Baker, M. S., Marko-Varga, G. & Rodriguez, H. Analytical validation considerations of multiplex mass-spectrometry-based proteomic platforms for measuring protein biomarkers. J Proteome Res 13, 5325–5332 (2014). 10.1021/pr500753r

11 Khamis, M. M., Adamko, D. J. & El-Aneed, A. Strategies and Challenges in Method Development and Validation for the Absolute Quantification of Endogenous Biomarker Metabolites Using Liquid Chromatography-Tandem Mass Spectrometry. Mass Spectrom Rev 40, 31–52 (2021). 10.1002/mas.21607

12 Doerr, A. DIA mass spectrometry. Nature Methods 12, 35–35 (2015). 10.1038/nmeth.3234

13 Bateman, N. W. et al. Maximizing peptide identification events in proteomic workflows using data-dependent acquisition (DDA). Mol Cell Proteomics 13, 329–338 (2014). 10.1074/mcp.M112.026500

14 Gillet, L. C. et al. Targeted data extraction of the MS/MS spectra generated by data-independent acquisition: a new concept for consistent and accurate proteome analysis. Mol Cell Proteomics 11, 0111 016717 (2012). 10.1074/mcp.O111.016717

15 Egertson, J. D. et al. Multiplexed MS/MS for improved data-independent acquisition. Nat Methods 10, 744–746 (2013). 10.1038/nmeth.2528

16 Gupta, S., Sing, J. C. & Rost, H. L. Achieving quantitative reproducibility in label-free multisite DIA experiments through multirun alignment. Commun Biol 6, 1101 (2023). 10.1038/s42003-023-05437-2

17 Anderson, L. & Hunter, C. L. Quantitative mass spectrometric multiple reaction monitoring assays for major plasma proteins. Mol Cell Proteomics 5, 573–588 (2006). 10.1074/mcp.M500331-MCP200

18 Kontostathi, G. et al. Development and Validation of Multiple Reaction Monitoring (MRM) Assays for Clinical Applications. Methods Mol Biol 1959, 205–223 (2019). 10.1007/978-1-4939-9164-8_14

19 Sandow, J. J. et al. Discovery and Validation of Novel Protein Biomarkers in Ovarian Cancer Patient Urine. Proteomics Clin Appl 12, e1700135 (2018). 10.1002/prca.201700135

20 Boja, E. S. & Rodriguez, H. Mass spectrometry-based targeted quantitative proteomics: achieving sensitive and reproducible detection of proteins. Proteomics 12, 1093–1110 (2012). 10.1002/pmic.201100387

21 Carr, S. A. et al. Targeted peptide measurements in biology and medicine: best practices for mass spectrometry-based assay development using a fit-for-purpose approach. Mol Cell Proteomics 13, 907–917 (2014). 10.1074/mcp.M113.036095

22 Hoofnagle, A. N. et al. Recommendations for the Generation, Quantification, Storage, and Handling of Peptides Used for Mass Spectrometry-Based Assays. Clin Chem 62, 48–69 (2016). 10.1373/clinchem.2015.250563

23 Grant, R. P. & Hoofnagle, A. N. From lost in translation to paradise found: enabling protein biomarker method transfer by mass spectrometry. Clin Chem 60, 941–944 (2014). 10.1373/clinchem.2014.224840

24 Grote, E., Fu, Q., Ji, W., Liu, X. & Van Eyk, J. E. Using pure protein to build a multiple reaction monitoring mass spectrometry assay for targeted detection and quantitation. Methods Mol Biol 1005, 199–213 (2013). 10.1007/978-1-62703-386-2_16

25 Shi, T. et al. Advances in targeted proteomics and applications to biomedical research. Proteomics 16, 2160–2182 (2016). 10.1002/pmic.201500449

26 Reker, D. & Malmstrom, L. Bioinformatic challenges in targeted proteomics. J Proteome Res 11, 4393–4402 (2012). 10.1021/pr300276f

27 Stewart, H. I. et al. Parallelized Acquisition of Orbitrap and Astral Analyzers Enables High-Throughput Quantitative Analysis. Anal Chem 95, 15656–15664 (2023). 10.1021/acs.analchem.3c02856

28 Heil, L. R. et al. Evaluating the Performance of the Astral Mass Analyzer for Quantitative Proteomics Using Data-Independent Acquisition. J Proteome Res 22, 3290–3300 (2023). 10.1021/acs.jproteome.3c00357

29 Guzman, U. H. et al. Ultra-fast label-free quantification and comprehensive proteome coverage with narrow-window data-independent acquisition. Nat Biotechnol (2024). 10.1038/s41587-023-02099-7

30 Baumgart, D. C. & Carding, S. R. Inflammatory bowel disease: cause and immunobiology. Lancet 369, 1627–1640 (2007). 10.1016/S0140-6736(07)60750-8

31 Strober, W., Fuss, I. & Mannon, P. The fundamental basis of inflammatory bowel disease. J Clin Invest 117, 514–521 (2007). 10.1172/JCI30587

32 Shanahan, F. Inflammatory bowel disease: immunodiagnostics, immunotherapeutics, and ecotherapeutics. Gastroenterology 120, 622–635 (2001). 10.1053/gast.2001.22122

33 Fakhoury, M., Negrulj, R., Mooranian, A. & Al-Salami, H. Inflammatory bowel disease: clinical aspects and treatments. J Inflamm Res 7, 113–120 (2014). 10.2147/JIR.S65979

34 Iskandar, H. N. & Ciorba, M. A. Biomarkers in inflammatory bowel disease: current practices and recent advances. Transl Res 159, 313–325 (2012). 10.1016/j.trsl.2012.01.001

35 Nowak, J. K., Kalla, R. & Satsangi, J. Current and emerging biomarkers for ulcerative colitis. Expert Rev Mol Diagn 23, 1107–1119 (2023). 10.1080/14737159.2023.2279611

36 Mc Ardle, A. et al. Standardized Workflow for Precise Mid- and High-Throughput Proteomics of Blood Biofluids. Clin Chem 68, 450–460 (2022). 10.1093/clinchem/hvab202

37 Fu, Q. et al. Highly Reproducible Automated Proteomics Sample Preparation Workflow for Quantitative Mass Spectrometry. J Proteome Res 17, 420–428 (2018). 10.1021/acs.jproteome.7b00623

38 Sakurai, T. & Saruta, M.Positioning and Usefulness of Biomarkers in Inflammatory Bowel Disease. Digestion 104, 30–41 (2023). 10.1159/000527846

39 Holzinger, D. & Foll, D. [Biomarkers for chronic inflammatory diseases]. Z Rheumatol 74, 887–896; quiz 897 (2015). 10.1007/s00393-015-0009-7

40 Mesaros, C. & Blair, I. A. Mass spectrometry-based approaches to targeted quantitative proteomics in cardiovascular disease. Clin Proteomics 13, 20 (2016). 10.1186/s12014-016-9121-1

41 Williams, D. K. & Muddiman, D. C. Absolute quantification of C-reactive protein in human plasma derived from patients with epithelial ovarian cancer utilizing protein cleavage isotope dilution mass spectrometry. J Proteome Res 8, 1085–1090 (2009). 10.1021/pr800922p

42 Miranda-Garcia, P., Chaparro, M. & Gisbert, J. P. Correlation between serological biomarkers and endoscopic activity in patients with inflammatory bowel disease. Gastroenterol Hepatol 39, 508–515 (2016). 10.1016/j.gastrohep.2016.01.015

43 Ricci, G., D’Ambrosi, A., Resca, D., Masotti, M. & Alvisi, V. Comparison of serum total sialic acid, C-reactive protein, alpha 1-acid glycoprotein and beta 2-microglobulin in patients with non-malignant bowel diseases. Biomed Pharmacother 49, 259–262 (1995). 10.1016/0753-3322(96)82632-1

44 Henry, O., Jourdan, B. & Duviquet, M. [Analysis of the inflammatory response in elderly hospitalized patients]. Ann Med Interne (Paris) 150, 189–194 (1999).

45 Fischer, K. et al. Biomarker profiling by nuclear magnetic resonance spectroscopy for the prediction of all-cause mortality: an observational study of 17,345 persons. PLoS Med 11, e1001606 (2014). 10.1371/journal.pmed.1001606

46 Fu, Q. et al. An Empirical Approach to Signature Peptide Choice for Selected Reaction Monitoring: Quantification of Uromodulin in Urine. Clin Chem 62, 198–207 (2016). 10.1373/clinchem.2015.242495

47 Sundararaman, N. et al. BIRCH: An Automated Workflow for Evaluation, Correction, and Visualization of Batch Effect in Bottom-Up Mass Spectrometry-Based Proteomics Data. J Proteome Res 22, 471–481 (2023). 10.1021/acs.jproteome.2c00671

48 Ebinger, J. E. et al. Seroprevalence of antibodies to SARS-CoV-2 in healthcare workers: a cross-sectional study. BMJ Open 11, e043584 (2021). 10.1136/bmjopen-2020-043584

49 Li, D. et al. The T-cell clonal response to SARS-CoV-2 vaccination in inflammatory bowel disease patients is augmented by anti-TNF therapy and often deficient in antibody-responders. medRxiv (2021). 10.1101/2021.12.08.21267444

50 Mujukian, A. et al. Postvaccination Symptoms After SARS-CoV-2 mRNA Vaccination Among Patients With Inflammatory Bowel Disease: A Prospective, Comparative Study. Inflamm Bowel Dis (2023). 10.1093/ibd/izad114

51 Li, D. et al. Post-Vaccination Symptoms after A Third Dose of mRNA SARS-CoV-2 Vaccination in Patients with Inflammatory Bowel Disease. medRxiv (2021). 10.1101/2021.12.05.21266089

52 Huang, Y. et al. Evidence of premature lymphocyte aging in people with low anti-spike antibody levels after BNT162b2 vaccination. iScience 25, 105209 (2022). 10.1016/j.isci.2022.105209

53 Fu, Q. et al. A Plasma Sample Preparation for Mass Spectrometry using an Automated Workstation. J Vis Exp (2020). 10.3791/59842

54 Demichev, V., Messner, C. B., Vernardis, S. I., Lilley, K. S. & Ralser, M. DIA-NN: neural networks and interference correction enable deep proteome coverage in high throughput. Nat Methods 17, 41–44 (2020). 10.1038/s41592-019-0638-x

